# Temporal scaling of ageing as an adaptive strategy of *Escherichia coli*

**DOI:** 10.1101/376566

**Authors:** Yifan Yang, Ana L. Santos, Luping Xu, Chantal Lotton, François Taddei, Ariel B. Lindner

## Abstract

Natural selection has long been hypothesised to shape ageing patterns, but whether and how ageing contributes to life-history evolution remains elusive. The complexity of various ageing-associated molecular mechanisms and their inherent stochasticity hinder reductionist approaches to the understanding of functional senescence, *i.e*. reduced fecundity and increased mortality. Recent bio-demographic work demonstrated that high-precision statistics of life-history traits such as mortality rates could be used phenomenologically to understand the ageing process. We adopted this approach to study cellular senescence in growth-arrested *E. coli* cells, where damages to functional macromolecules are no longer diluted by fast *de novo* biosynthesis. We acquired high-quality longitudinal physiological and life history data of large environmentally controlled clonal *E. coli* populations at single-cell resolution, using custom-designed microfluidic devices coupled to time-lapse microscopy. We show that *E. coli* lifespan distributions follow the Gompertz law of mortality, a century-old actuarial observation of human populations, despite developmental, cellular and genetic differences between bacteria and metazoan organisms. Measuring the shape of the hazard functions allowed us to disentangle quantitatively the demographic effects of ageing, which accumulate with time, from age-independent genetic longevity-modulating interventions. A pathway controlling cellular maintenance, the general stress response, not only promotes longevity but also temporally scales the whole distribution by reducing ageing rate. We further show that *E. coli*, constrained by the amount of total biosynthesis, adapt to their natural feast-or-famine lifestyle by modulating the amount of maintenance investment, rendering ageing rate a highly evolvable life-history trait.

## Background

The biology of ageing and senescence is centred on the duality of individual frailty and population resiliency. Throughout the course of normal metabolism, components of living systems such as cells, lipids, proteins and DNA inevitably suffer from wear-and-tear such as free-radical damages. This constant and collective decay eventually leads to the loss of vital functions and collapse of individuals. Understanding the way that system failure emerges out of distributed microscopic damages could reveal how functional components are organised into self-maintaining individuals in the first place^1,2^. Yet for some organisms in the tree of life, ageing does not lead to increased mortality and declined fertility^3^. Organisms possess the abilities to repair or replace most of the damages to their components, exemplified by the “immortal germ line”^4^. Evolutionary biologists attribute the apparent senescence of metazoan somas to the inadequate investment in cellular maintenance as an adaptive strategy to maximise lifetime reproductive success, due to the trade-offs between the survival and reproduction of the young on one hand, and maintenance for the benefit of the old on the other^5,6^. Presumably, natural selection has to operate through the molecular “levers” of damage accumulation and/or repair to achieve such life-history optimisation.

*Escherichia coli*, a single-cell prokaryote with short lifespans, has historically served as a model organism that resolved many fundamental questions in biology. *E.coli* cells, as their metazoan counterparts, suffer from damages to their components, which lead to cellular senescence^7,8^. In exponential growth, these damages are quickly diluted by *de novo* biosynthesis, and the effects of cellular senescence mitigated by rapid and robust reproduction^9^. Yet, the natural life cycle of *E.coli* entails a much wider range of physiological conditions than exponential growth. Most of bacterial cells spend much of their lives in resource-limited growth arrested conditions, where *de novo* biosynthesis rates are slower^10^ and cells undergo senescence due to the accumulation of molecular damages such as protein misfolding and oxidation (Fig. 1)^11^. Despite the lack of fixed separation of germ and soma cells, the ability to survive the wear and tear of cellular components during growth arrests contribute to bacterial overall fitness as much as the ability for exponential growth. We, therefore, adopted a bio-demographic approach^1,2^ to understand how modulation of molecular damage repair could shape ageing dynamics in growth-arrested *E.coli*.

**Figure 1.**
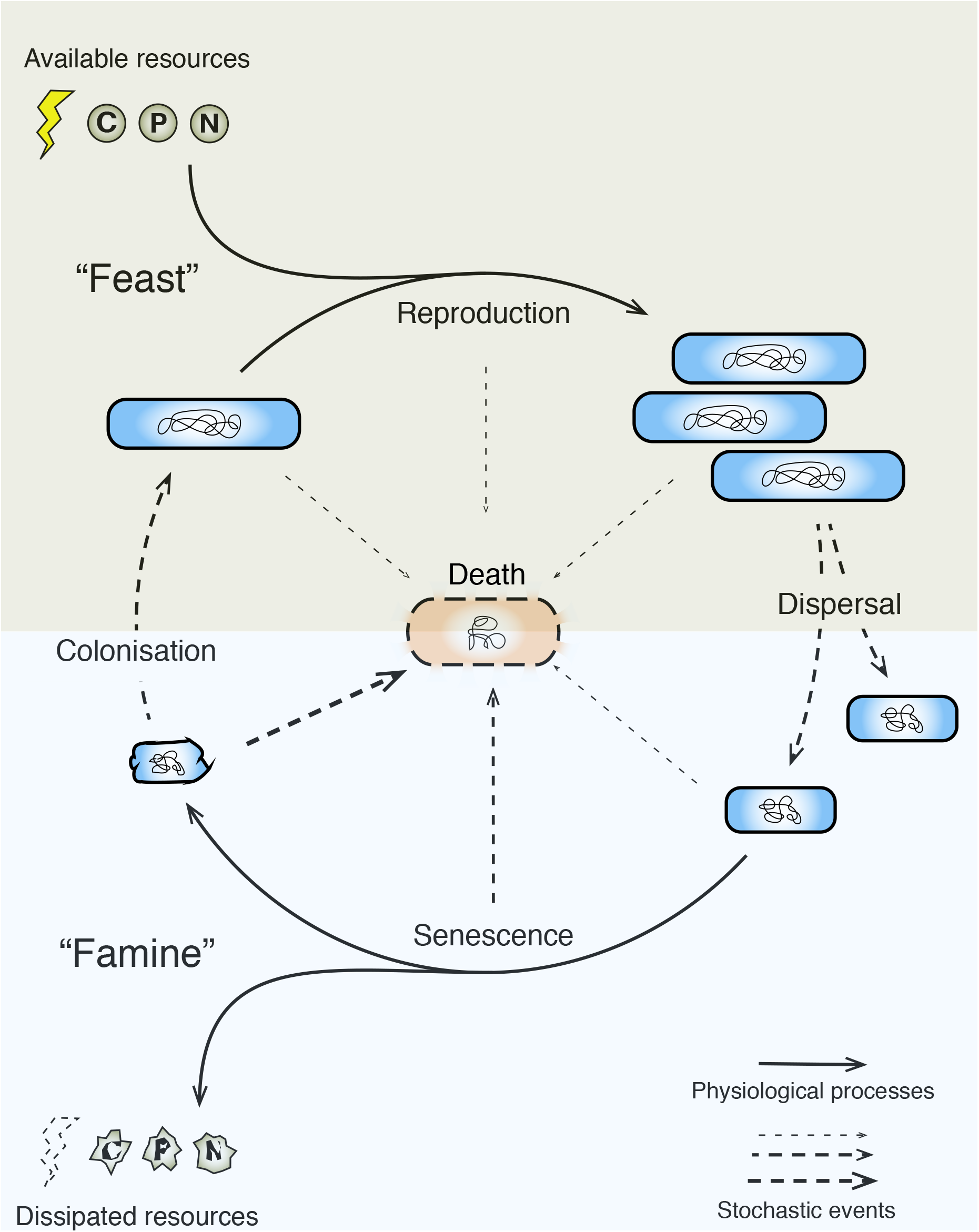
The feast-and-famine life cycle of *E. coli*. Single-cell organisms such as *E. coli* undergo developmental transitions not in response to fixed developmental programs but opportunistically in response to environmental changes. Because long-term growth rates in a stable ecosystem remain close to zero, life-history traits in “feast” and “famine” conditions should contribute rather equally to overall fitness. In “feast” conditions, the traits under strong selection are resource assimilation and reproduction, while in “famine” conditions are maintenance and survival. Senescence has been shown to occur in “famine” conditions^11^.

### Custom-designed microfluidic devices allow longitudinal tracking of vitality at single-cell level

We first set out to acquire high quality longitudinal physiological and life-history data at single-cell resolution of clonal populations of *E.coli*. For decades, limitations of traditional cell culture methods have frustrated quantitative physiologists’ attempts to understand bacterial maintenance^12,13^, due to both media-cell interactions and cellular interaction such as cross-feeding and cannibalism. To be able to measure individual *E.coli* cell lifespan in constant and homogenous environmental settings, we designed a novel microfluidic device with cell-dimension chambers (Fig. 2). Cells are trapped in an array of single-cell-wide dead-ended wells, with openings to a main flow channel. Constant flow of fresh media in this main channel provides necessary nutrients and eliminates metabolic waste, cell debris and intercellular crosstalk so that the environmental conditions are maintained constant over time. Cells with appropriately expressed fluorescent markers are imaged bottom-up and appear as fluorescent spots (Fig. 2b). This experimental system allows easy tracking of cohorts of a large number of individual bacterial cells for prolonged periods (up to 7 days).

**Figure 2.**
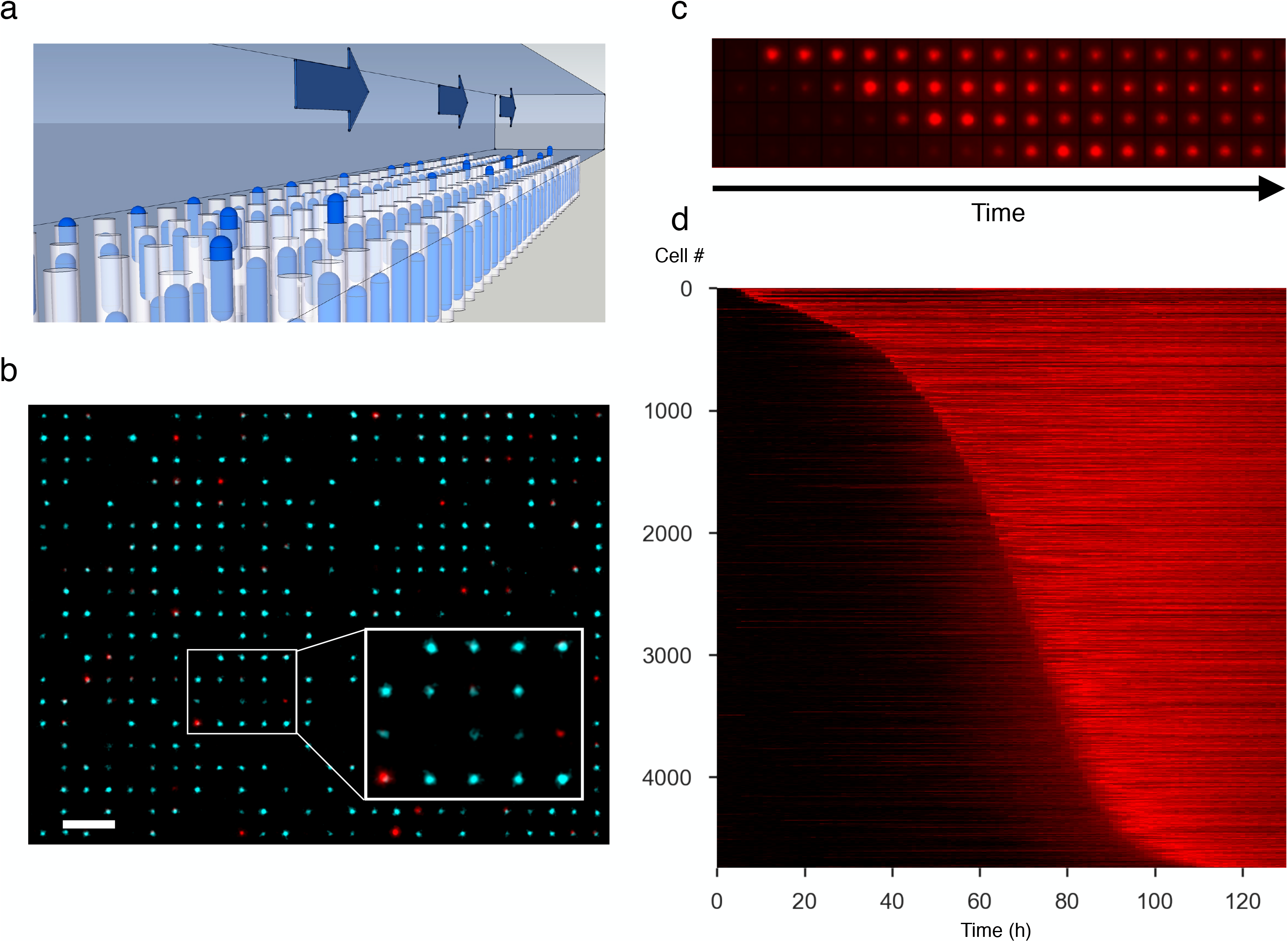
Coupling microfluidic chip and time-lapse microscopy to measure *E. coli* lifespan distribution at single-cell resolution. **a**, 3D model of microfluidic devices used to trap and isolate large number of single cells. The blue-gray cube in the upper half of the picture represents the main flow channel where fresh, carbon-source-free media are supplied. The array of light-blue rods represents *E.coli* single-cells trapped in a 2D-array of cell-sized dead-end chambers. The arrows represent the media flow that maintain the environmental homogeneity and removes debris. **b**, Fluorescence microscopy image of the microfluidic device loaded with *E.coli* cells. Each fluorescent dot corresponds to a single cell trapped in the 2D array of dead-end wells. Z-axis focus is adjusted so that dead-ends of the micro-wells are imaged. The cyan pseudo-colour represents constitutively expressed fluorescent protein fluorescence signal; red pseudo-colour - Propidium Iodide (P.I.) fluorescent signal described in the text. **c**, Sample time-lapse P.I. images of mortality events, from early (top) to late (bottom) deaths. **d**, Heat map of P.I. signal time-series for a population of single cells. The colour values of each row correspond to the time-series of one cell, ranked according to lifespan, top-to-bottom, from the shortest living to the longest living cell. The boundary between dark and bright red indicates the transition from life to death and defines the lifespan distribution. Dark strips after cell deaths indicate decaying DNA or empty wells after cell debris have been washed away. Data shown in **d** have been smoothed according to the procedure described in supplementary material to remove minor signal fluctuations coming from microscopy focusing. The same procedure is used before time-of-death is estimated.

In order to observe cell mortality, we included in the carbon-source-free medium a red-fluorescent, DNA-binding, bacterial viability dye, Propidium Iodide (P.I.), which penetrates the cells only when cellular membrane potentials are disrupted. P.I. staining has been established as an effective proxy of cellular death and correlates well with cell viability assessed by proliferating potential^14^. We used P.I. at a concentration 4-fold lower than the previous concentration that had no effects on *E.coli* viability and growth^14^. Automatic time-lapse fluorescence microscopy and fixed geometry of our devices allowed longitudinal quantification of P.I. signal for every single cell in the population. We defined the half point between peak P.I. signal and background fluorescence as a threshold to establish the time-of-death. At the end of the experiments, 70% ethanol was injected into the device in order to account for every cell in the cohort, and establish a lower bound for the time-of-death of surviving cells. These cells were censored at the time that their P.I. signals crossed their respective thresholds (see Methods).

Despite being genetically clonal and environmentally controlled, cells did not share the same time or manner of death, as measured by the P.I. time-series. The transition from life to death of a representative cohort (N= 4744) could be visualised by the boundary between dark to light in Fig. 2d. Because cells were sorted vertically according to estimated lifespans, the shape of this boundary represents the survival function of the population. Considering ageing and death as a stochastic process of system reliability reduction^1,2^, the observed time-of-death distribution could be viewed as the population’s first-passage time distribution. In addition, we observed individual differences in death trajectories. Short-lived cells tended to have very sharp P.I. increases that associated with abrupt losses of membrane integrity; while those that die late tended to suffer a type of “slow death” characterised by a gradual P.I. increase over the course of 10-15 hours.

### *E. coli* lifespan distributions follow the Gompertz law

For both demographers and reliability engineers, the age-associated increase in death probability, also known as the hazard rate *h*(*t*), is considered to be a hallmark of ageing^1–3^. The detailed population statistics derived from our experiments are particularly useful in estimating the shape of the hazard functions over the whole lifespan. We find that *E.coli* lifespan distributions have, as their main feature, regimes with exponentially increasing hazard rates, *i.e*. the Gompertz law of mortality *h*(*t*) ~ *h*(*t_0_*)*e*^*b*(*t*−*t0*)^, where *h*(*t_0_*) is the initial hazard rate and *b* is the Gompertz ageing rate^15,16^. We estimated *h*(*t*) directly with binomial error by binning mortality events within discrete time intervals (τ_i_, τ_i+1_), *i.e*. 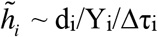, where Y_i_ is the number of individuals at risk at time τ_i_ and d_i_ is the number of cell deaths within (τ_i_, τ_i+1_) (see Fig. 3). For wildtype cells, this exponential regime spans from about 13 hours to 93 hours, corresponding to approximately 90% of all cell deaths (N_total_=4744, see Fig. 3b, top x-axis), and ranges at least 100-fold changes in hazard rates (2*10^−3^ h^−1^ to 2*10^−1^ h^−1^). Within this exponential regime, the doubling time of hazard rate is 9.4±0.5 hours. In comparison, the doubling time of human mortality hazards is about 8 years^15^.

**Figure 3.**
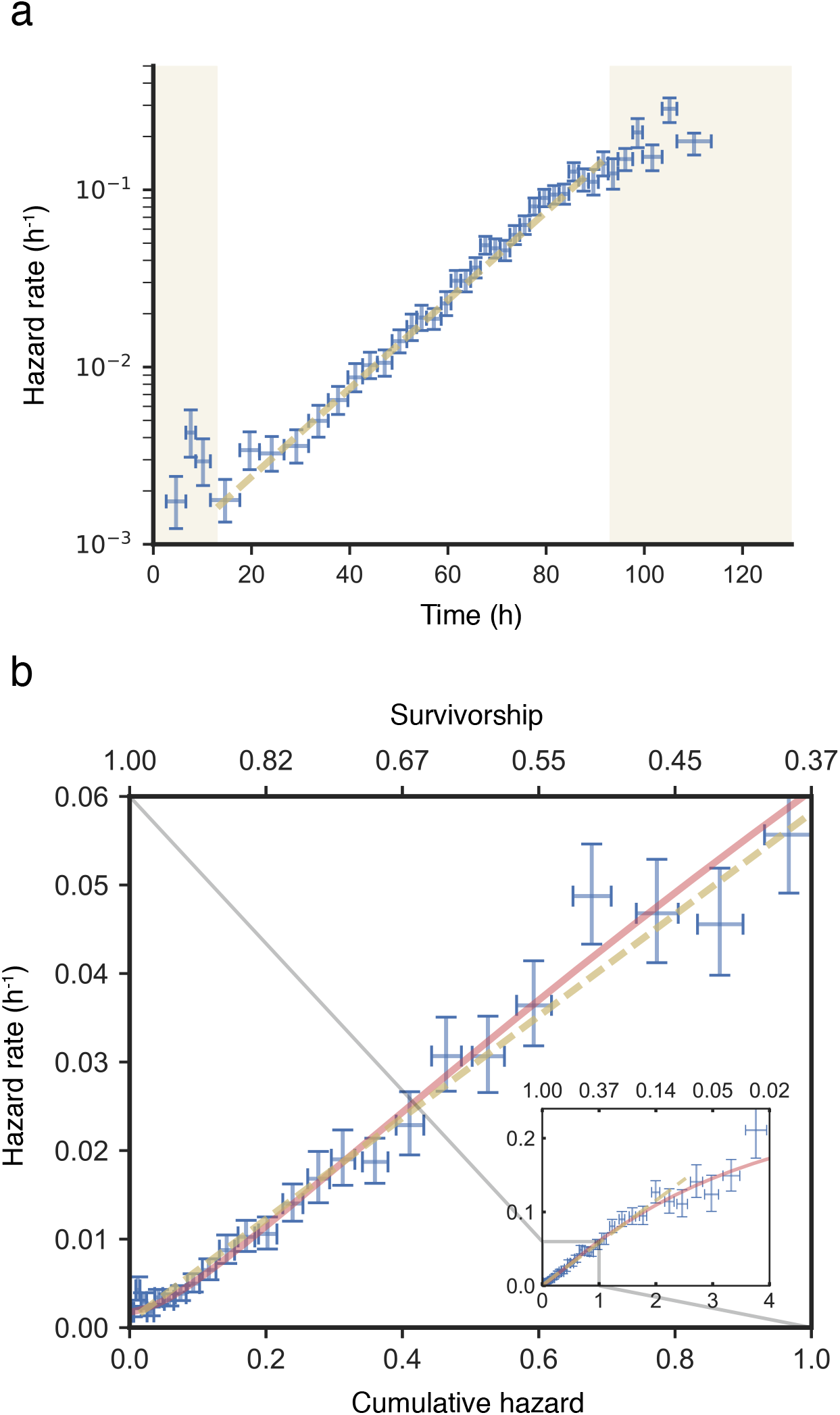
Non-parametric estimation indicates that wildtype *E. coli* lifespan distribution can be characterised by the Gompertz law. Shown are non-parametric estimations of the hazard functions and cumulative hazard functions of the same populations in one particular microfluidic experiment. **a**, Binomial estimators of the hazard rates, shown in log scale. Cell deaths are binned into discrete time intervals, which are marked by the x-axis error bars. The time bins are chosen so that the Nelson-Aalen confidence intervals (CI) in b do not overlap with each other. Error bars on the y-axis are the 95% CI for hazard rate estimated based on binomial distribution. Yellow shading covers regions that deviate from the exponential hazard regime for the shortest and longest living individuals. **b**, Cumulative hazard dynamics in the phase plane. Data, binning intervals and y-axis are the same as those in **a**. Instantaneous hazard rates plotted against the cumulative hazards (bottom x-axis), or equivalently the negative logarithm of survivorship (top x-axis). Nelson-Aalen estimators are used for the cumulative hazard function. The horizontal error bars are the 95% CI of the Nelson-Aalen estimator at the centre of the time bins. Dashed lines are maximum likelihood parametric estimations using the Gamma-Gompertz-Makeham model. The inset provides a zoom-out view of the whole data range, while the main figure zooms in on the first 63% of cell deaths.

Hazard rates not only define lifespan distributions but can also be thought of as surrogates for system vulnerability, whose dynamics through time reflect the physiological consequence of ageing. In this light, the Gompertz law can be interpreted as a dynamic equation governing the ageing process: *dh*(*t*)/*dt* = *bh*(*t*), where *b* is the Gompertz ageing rate.

The integral version, *h*(*t*) = *bH*(*t*)+*h*(*t_0_*), where 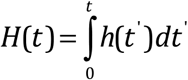, can be directly observed in the phase plane of the cumulative hazards *H*(*t*) without any free parameters (Fig. 3b). The integral equation is used because *H*(*t*) can be estimated independent of binning, by using the Nelson-Aalen (N-A) estimator 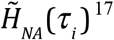^17^. The fact that the theoretical trajectory, including the aforementioned linear regime, lies within the confidence intervals (CI) of almost every state coordinate 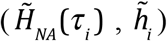 lends strong statistical credence to the Gompertz law for describing our data. In addition, the observed trajectory in the phase plane links the instantaneous hazard rate *h*(*t*) at any given time *t*, with the total proportion of population that have died before time *t* (because *H*(*t*) = −*ln*[*S*(*t*)], where *S*(*t*) is the survival function).

The hazard rate observed deviates from the exponential regime at the beginning (t<15 hours, 3% cell deaths) and the end (t>93 hours, 7% cell deaths) of the total lifespan. These observations are reminiscent of similar deviations from Gompertz law in human mortality data^18,19^. In our case, the additional mortality at the early age might result from harvesting and transferring exponentially growing cells from batch culture directly to growth-arresting conditions inside our microfluidic chip, in a way analogous to infant mortality in human mortality data, which can be modelled by an age-independent component (*λ* in the Gompertz-Makeham model 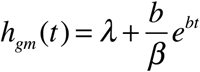, where *β* controls the age at which the Gompertz regime overtakes the Makeham term *λ*).

Late-life hazard decelerations are common phenomena for both model organisms and human populations^18,20^. It is thought that these hazard decelerations often do not reflect ageing decelerations for individuals, but result from changes in composition of heterogeneous populations. The more fragile individuals are more likely to die and be removed from the cohort. When hazard rates are high late in life, significant portions of the populations are removed so that the observed hazard rates of the surviving populations would be smaller than the original cohorts on average. We found that an extension to the Gompertz law named Gamma-Gompertz, originally used to model old-age human model data^21^, could also satisfactorily model late life decelerations in our data, for both wildtype and mutant strains (see below). It could be understood as accounting for the compositional changes by an additional parameter, s, controlling the level of frailty heterogeneity: 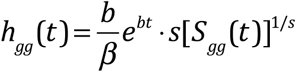, where S_gg_(t) is the survival function of Gamma-Gompertz, and other parameters are as before. Despite being genetically and environmentally non-distinguishable, we think it is reasonable to assume frailty heterogeneity exist in our single-cell populations due to the stochastic nature of cellular biochemistry, explaining the suitability of the Gamma-Gompertz for our data.

We thus used the Gamma-Gompertz-Makeham (GGM) extension model with 4 parameters to fully model our data across the whole lifespan, with parameters estimated by likelihood maximisation:

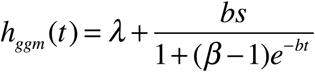

Goodness-of-fit was evaluated using a one-sample Kolmogorov-Smirnov test for right-censored survival time data. The appropriateness of the model terms was evaluated by comparing Akaike Information Criterion (AIC) of the candidate models (see methods).

### The general stress response of *E. coli* modulates the ageing rate

Evolutionary theories of ageing predict the existence of ageing-modulating mechanisms and their activation by nutrient limitation^22^. Several nutrient-sensing pathways in both bacteria and metazoans have been shown to control stress resistance and reduce mortality^23–25^. Yet their roles in delaying ageing-related damage accumulations are often controversial. It is often difficult and requires large experimental cohorts to disentangle their effects on ageing from age-independent components of longevity^26^. Having established a method to measure ageing rates using high-quality mortality statistics, we harnessed this system to shed light on potential ageing-modulating mechanisms of *E.coli* that might be activated in response to nutrient deprivation.

Many bacteria species regulate the level of cellular maintenance through a genetic pathway called general stress response (GSR), controlled by the master transcriptional regulator *rpoS* that is activated by nutrient-deprivation among other signals^27^ (Fig. 4a). To assess the role of rpoS-controlled cellular maintenance in the ageing of growth-arrested *E.coli*, we measured the lifespan distributions of two GSR mutants, Δ*rpoS*, and Δ*rssB* with that of the wildtype strain. Δ*rpoS* is the null mutant and Δ*rssB* displays an elevated GSR due to increased RpoS stability^28^. We observed that higher GSR promotes longevity in the microfluidic experiments, whereas the absence of GSR results in shortened longevity (Fig. 4b and Fig. 5a). Our large sample sizes allowed us to directly measure the hazard dynamics (Fig. 4c) of each strain, which could disentangle GSR’s effects on ageing rate, as opposed to age-independent components of longevity.

**Figure 4.**
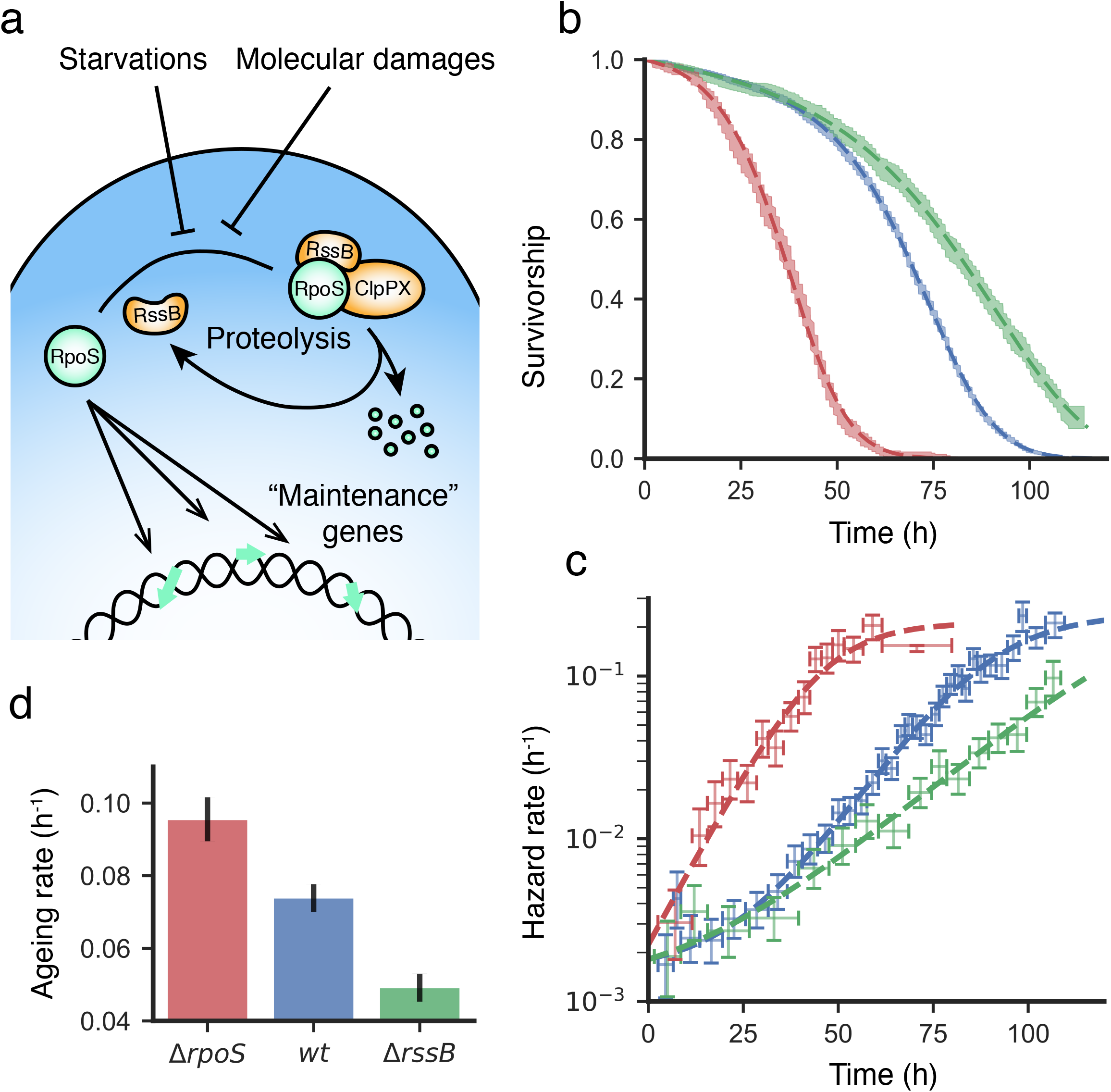
The general stress response of *E. coli* modulates ageing rate. The lifespan distribution for the wildtype (*wt*), Δ*rpoS* (lacking general stress response) and Δ*rssB* (overexpressing the general stress response) strains are measured multiple times by independent microfluidic experiments. **a**, A scheme representing relevant regulatory features of the general stress response, and in particular the functions of the genes *rpoS* and *rssB*. **b**, Experimental and GGM model survivorship. Representing the experimental survivorship, colour bands are the 95% CI of the Kaplan-Meier (K-M) estimators. Coloured dashed lines are Gamma-Gompertz-Makeham (GGM) models whose parameters are estimated from maximum-likelihood (ML) methods. **c**, Hazard rates estimated using only cell deaths within discrete time intervals (Error bar markers), and GGM hazard models estimated from the whole dataset using ML methods. Similar to Fig. 3a, Vertical error bars are binomial 0.95 confidence intervals and horizontal error bars are the binning time intervals. Data from b and c are from the same representative experimental cohorts for each strain. **d**, Ageing rates for each strain, estimated by ML-fitted GGM models to 3 independent experiments. Error bars represent 95% CI. The ageing rates are: *b_rpoS_* = 0.095±0.006 h^−1^, *b_rssB_* = 0.049±0.004 h^−1^ and *b_wt_* = 0.074±0.004 h^−1^.

**Figure 5.**
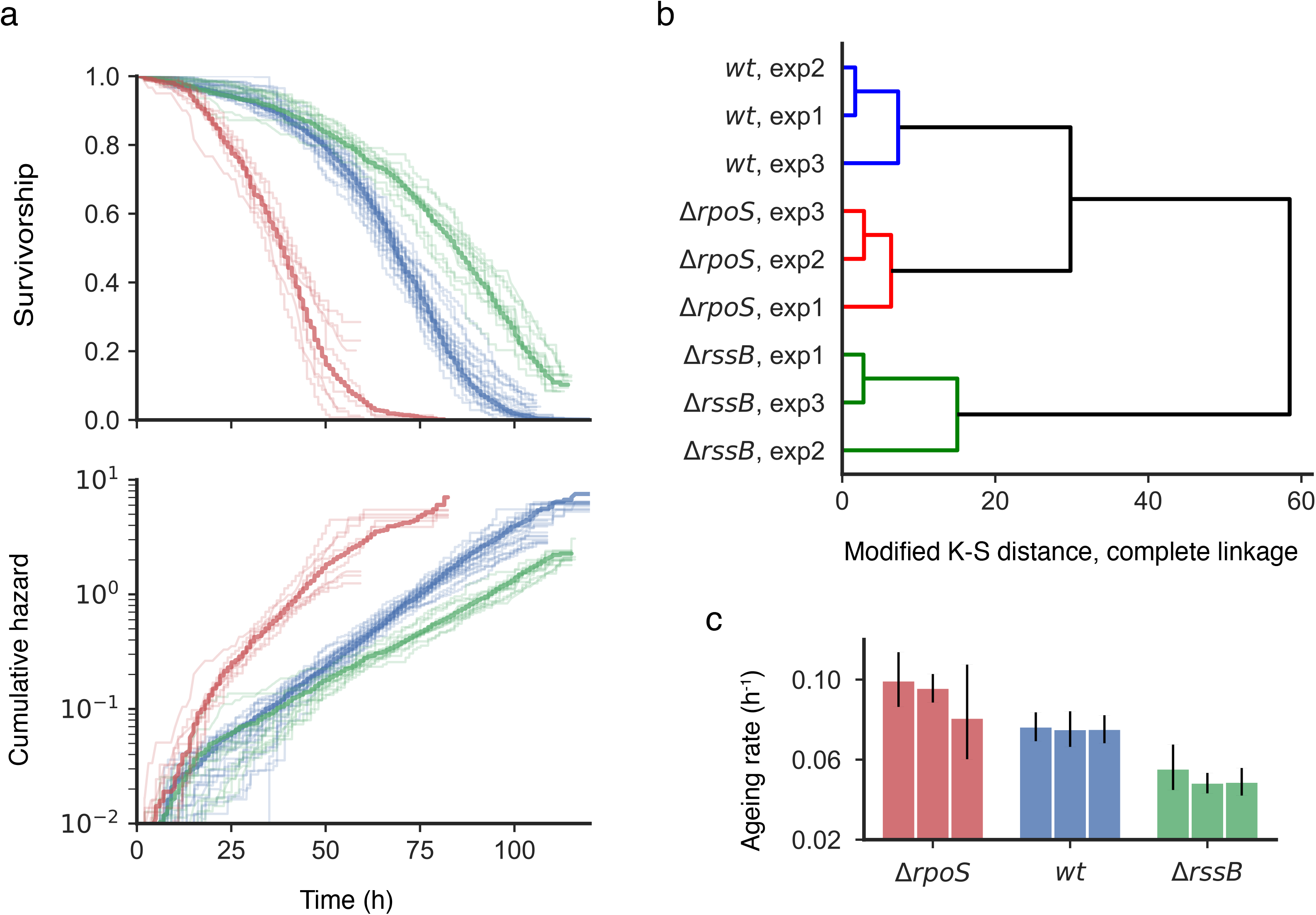
Experimental repeatability and variances. For each of wildtype (*wt*), Δ*rpoS* and Δ*rssB* knockout strains, data from 3 independent experimental replicates are shown. **a**, Survivorship (Kaplan-Meier estimator, K-M) and cumulative hazards (Nelson-Aalen estimator, N-A) of sub-populations from different imaging positions. Thick opaque lines correspond to the data shown in Fig. 4b and c. Thin semi-transparent lines are the K-M and N-A estimators of sub-populations from different imaging positions, which are used to constitute the population represented by the opaque lines. Each sub-population is from one imaging position that corresponds to a roughly 125μm-sided square-patch on the microfluidic chip, and consists of 150-600 cells. Imaging positions with less than 50 cells are not shown. **b**, Hierarchical clustering of the 3 independent experimental replicates of each strain. Standard agglomerative clustering algorithm is applied using complete linkage and the 2-sample modified Kolmogorov-Smirnov statistic sup{|Y(t)|} (See methods) as a distance metric. **c**, Ageing rates 3 independent experimental replicates of each strain. Ageing rates are estimated similarly as those in Fig. 4d, but independently for each experimental replicate without pooling them together using GLM. Error bars represent 95% CI.

Significantly, we found that increased GSR reduces the rate of ageing (Fig. 4d and Fig. 5c). Given how well the GGM model fitted for all 3 strains, the impact of genotypes on ageing parameters such as ageing rates could be extracted using generalised linear models (GLM). With enhanced GSR, Δ*rssB* cells double their mortality risk every 14.1 hours, with a 95% CI ranging from 13.1 to 15.3 hours, compared to 9.4 hours for the wildtype (CI 8.9 - 9.9 hours) and 7.3 hours for the null strain Δ*rpoS* (CI 6.8 to 7.7 hours). Variability of ageing rate measurements was assessed using 3 independent experimental cohorts for each strain (Figure S1). We visualised non-parametrically the overall experimental variations (Fig. 5a and b). Parametric differences in ageing rates among experimental repeats were tested using GLM models and AIC. We confirmed that experimental repeats shared similar ageing rates (H0: Same aging rate for experimental repeats. d.f. H_0_ - H_1_ = -2; ΔAIC_*rssB*_=-2.56, ΔAIC_*wt*_=-2.26, ΔAIC_*rpoS*_=-2.30; N_*wt*_=6867, N_*rssB*_=6969, N_*rpoS*_=4793).

The systematic increase in vulnerability of *E.coli* in our experiments is likely driven by the catabolism of pre-existing macromolecules and dissimilation of biomass^29,30^, which is necessary to provide energy to express housekeeping genes and maintain physiological homeostasis^10,31^. Our finding that GSR modulates ageing rate suggests that optimising this maintenance energy requirement is likely one of the physiological functions of the RpoS regulon (see Discussion).

### Evolutionary trade-offs mediated by the general stress response

Is slower ageing an adaptive life-history trait? This was one of original question raised by Sir Peter Medawar^32^ that motivated much of later ageing research. Optimal life-history theory suggested that trade-offs and constraints among fitness components shape metazoan ageing rates^6^. The possibility of modulating ageing rate through GSR offers us the opportunity to test these ideas in a fast-evolving organism as *E. coli*.

The relatively well-understood GSR pathway provides a clear molecular mechanism for a trade-off between growth and maintenance. The master regulator *rpoS* encodes the RNA polymerase (RNAP) sigma subunit σ^S^, which competes with the other sigma factors including the vegetative σ^D^ to recruit the core RNAP and direct the transcription and translation machinery towards the RpoS regulon. By titrating protein synthesis activity away from metabolic and ribosomal genes controlled by σ^D^, RpoS activity inhibits growth and nutrient assimilation^33^. We measured quantitatively the growth impact of modulating GSR levels, and modelled its effect using a simple course-grained model of proteome sectors^34^. We found that the proportion of protein synthesis devoted to the RpoS regulon linearly increases the timescale of growth (see top axis in Fig. 6a).

**Figure 6.**
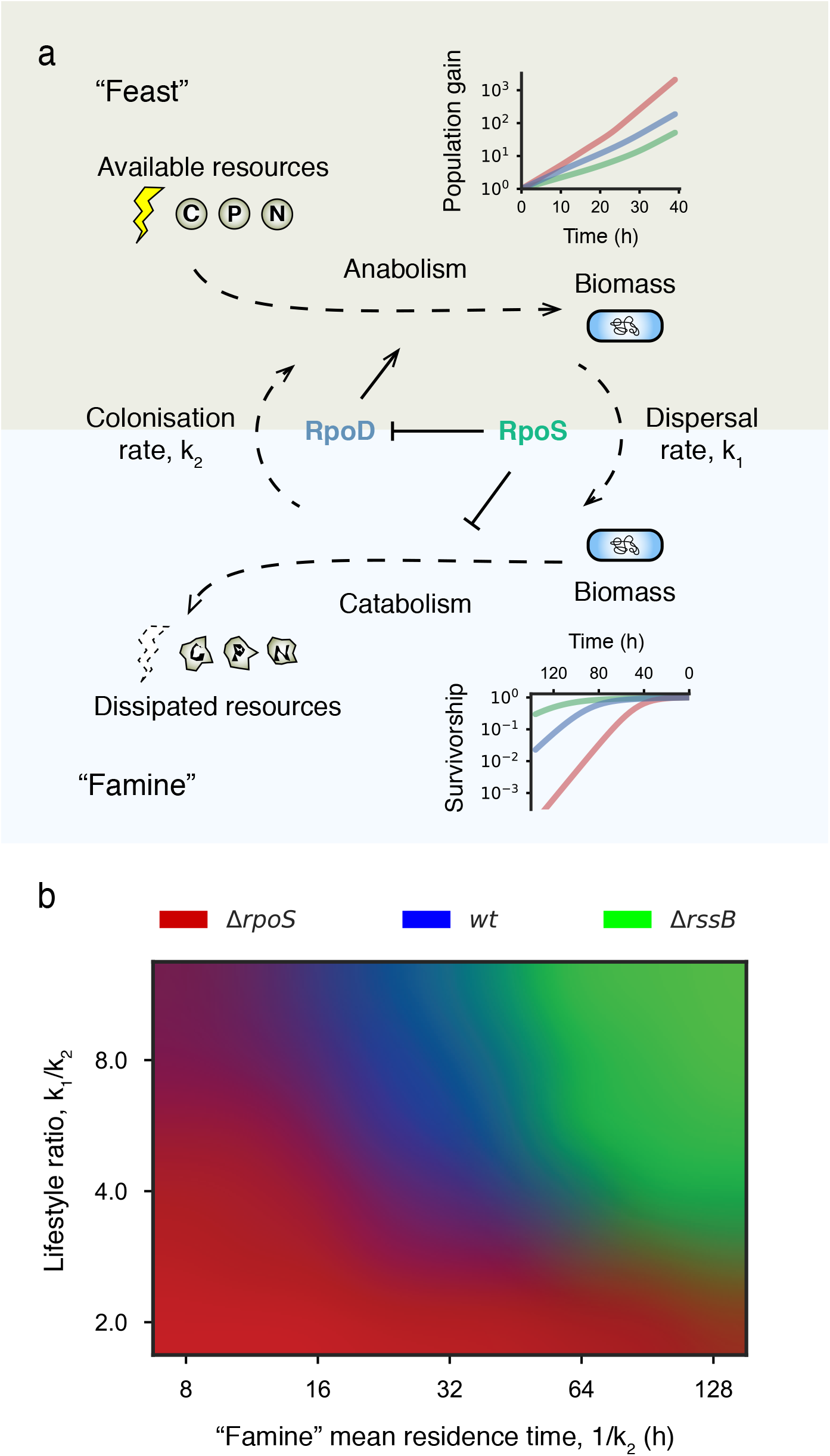
Trade-offs between growth and maintenance mediated by the *rpoS* pathway, and its fitness consequences. **a**, Scheme of the ecological processes (dashed arrows) and regulatory relationships (solid arrows) involved in the trade-offs mediated by *rpoS*. Environments are structured by alternating episodes of “feast” and “famine”. Fitness of Δ*rpoS* (red), *wild-type* (blue) and Δ*rssB* (green) as functions of time spent in each environmental episode (age) are plotted in the top and bottom axes. Fitness is defined as the logarithmic change of population sizes. **b**, Fitness comparison of the 3 strains across a range of environmental conditions to identify regimes favouring faster and slower ageing strategies. The colour-coded regions identify environmental conditions under which one strain dominates over the other two. Absolute fitness of the 3 strains are converted to RGB tuples using softmax normalisation. For each strain and each pair of environmental parameters, fitness is calculated by averaging the population growth/decline rates over 5000 episodes of both “feast” and “famine”. Episode lengths are independently drawn from exponential distributions, parameterised by the two ecological rates, k_1_ and k_2_, as shown in **a**. The fitness functions, population gain for “feast” and survivorship for “famine” (shown in the axes in **a**), are used to determine population size changes in each episode. See methods for details.

To assess the life-history optimality of different ageing rates, we integrated numerically the experimentally derived growth rates and mortality rates into fitness, defined as the long-term population growth rates. In contrast to metazoans, for whom fertility and mortality schedules are connected through fixed age structures, our model of *E. coli* life-history consisted of alternating environmental episodes of varying durations, in which *E. coli* populations either grow (“feast”), or decline (“famine”) (Fig. 6a). These feast-or-famine cycles were parameterised by two transition rates, controlling respectively the average lengths of “feast” and “famine” episodes. In order to identify selective pressure for ageing rates, we directly compared the fitness of the three strains with different GSR phenotypes, each representing a different strategic position in the growth-maintenance trade-off. We identified the environmental regimes selecting for faster or slower ageing (Fig. 6b). The boundaries between these regimes were characterised along two axes: lifestyle ratio, defined as the ratio of time spent in “famine” versus “feast”; and “famine” mean residence time, or in other words, average length of “famine” episodes.

There are two necessary conditions for selecting slower ageing strategies represented by Δ*rssB*. First, populations have to spend much more time in “famine” rather than “feast”, so that over the long term, population decline rather than grow. Secondly but no less important, given the same lifestyle ratio, famines should consist of longer episodes rather than short but more frequent ones. This condition is necessary due to the exponential mortality dynamics described by Gompertz law: investing in cellular maintenance only becomes beneficial at old age, when the exponentially increasing benefits of slower ageing eventually overcome the more immediate cost on growth. It is the typical timescale of “famine” that provides the selective pressure for ageing rates.

Despite representing the complex regulations of GSR with two mutants, we can now understand the ecological role of GSR activation and its adaptive consequences. Activated by declining nutrient availability, GSR directs resources towards internal maintenance to wait out the adverse conditions, whose lengths determine the optimal activation level. Previous observations from experimental evolution of *E. coli* support our predictions. In continuous cultures of *E. coli* where the populations do not pass through prolonged growth-arresting bottlenecks, mutations that attenuate or knock out RpoS activity are among the first to arise^35,36^. In contrast to the isolated populations under constant environmental conditions in our chip experiments, *E. coli* populations in nature influence their environments, and also interact with each other. These interactions may give rise to frequency-dependent selection and evolutionary game dynamics between slower and faster ageing strategies, as is observed in experimental evolution^37,38^.

## Discussion

We obtained high quality single-cell demographic data, using a novel microfluidic device, to demonstrate that ageing of growth-arrested *E. coli* follows the Gompertz law of mortality. Moreover, bacterial general stress response could temporally rescale the lifespan distribution by modulating the ageing rate. We further articulated in a demographic model the trade-offs and selective pressure driving the evolution of ageing rate.

In our work, two different conceptual perspectives of ageing are integrated and applied to one of the most iconic model organism, *E. coli*. One perspective, held by biochemists and physicists, sees ageing as the stochastic and inevitable erosion of organismal order and biochemical redundancy, created by self-reproducing and self-maintaining networks during growth and development. The other perspective, from evolutionary biology, views ageing as a component of the organismal life-history strategy, optimised and fine-tuned by natural selection. These two approaches constitute the proximate and ultimate causes of ageing respectively, with the former providing the constraints and the “lever” for the latter. In bacteria, we indeed observed the stochastic process of ageing and mortality. Lifespan vary significantly among genetically identical individuals in constant, homogeneous environment. However, at the population level, ageing is characterised qualitatively by the Gompertz law of mortality. In response to environmental conditions, the lifespan distribution is modulated by regulatory and selective forces through temporal scaling, while its general, exponential, shape is preserved. These new empirical observations provided an integrated perspectives to bacterial ageing, and by extension, potentially to the breath of the tree of life.

Although the Gompertz law has been shown to characterise ageing of many metazoan organisms, the physiological correlates of its parameters are not well understood. To rescale the lifespan distribution without changing the general shape of hazard dynamics, GSR has to orchestrate a coordinated response to manage various molecular damages that the organism encounters^1^. Indeed, RpoS regulates hundreds of genes conferring resistance to both internal and external stresses such as oxidative, thermal, acid, alkaline, osmotic, and UV^39^. For this reason, the ageing rate in our case reflects the general level of macromolecular catabolism and may relate to the maintenance energy of bacteria^13^. We hypothesise that the energetic costs of homeostasis and molecular damage repair are paid by the loss of biomass, leading to the gradual increase in the probability of death. It is the rate of energy dissipation, or the rate of living, that correlates with the ageing rate in our system (see Fig. 7).

**Figure 7.**
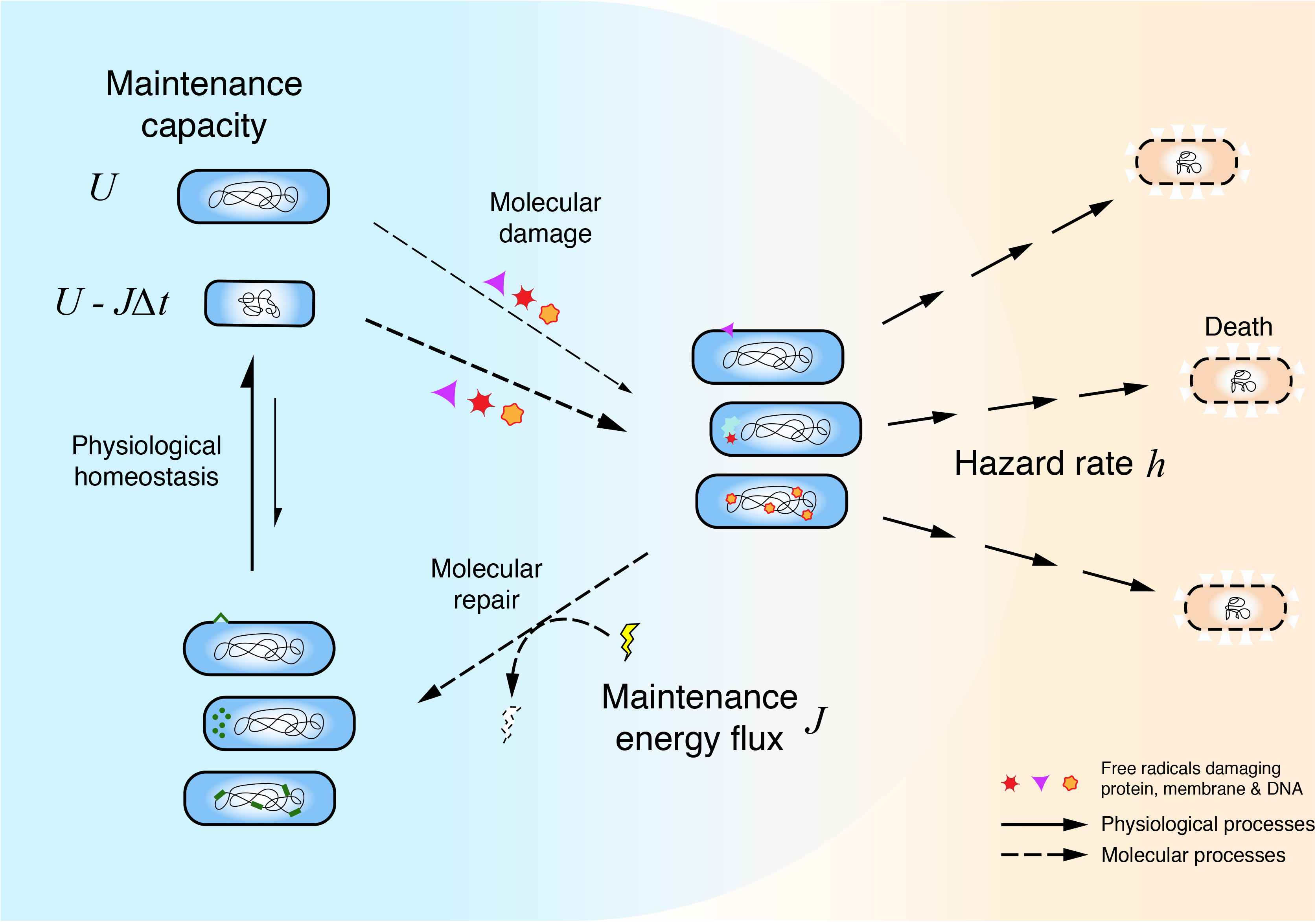
Gompertz law of mortality informs and constraints physiological models of ageing and mortality in growth-arrested *E.coli* cells. Depicted are the abstract states and pathways cells pass through during ageing and mortality. Acute damages to essential cellular components, when unrepaired, often lead to cell death, quantified by *h*(*t*). Cells could also triage and repair these damages to return themselves back to physiological homeostasis. The energy cost of this maintenance process, *J*(*t*), termed maintenance energy^13^, need to be repaid by either an external energy source or internal storage in the form of existing biomass. If we assume it is the same set of acute molecular damages that kill the cells or lead to ageing, then *J*(*t*) could relate to *h*(*t*) proportionally, *J*(*t*) = *Rh*(*t*) (eq1), where *R* is the “friction” coefficient controlled by the relative proportions of the two fates of damaged cells: repair vs death. For carbon-starved cells in our experiments, this flux of energy dissipation decreases the capacity for molecular repair in the future, *U*(*t*), leading to cellular senescence. Put it another way, *dU*(*t*)/*dt* = −*J*(*t*) (eq2). Through these assumptions, the hazard dynamics we described in this work could be recast as the dynamics of maintenance capacity. Our experimental results place non-trivial constraints on how the amount of maintenance capacity regulates the rate at which cells are damaged molecularly in the first place. If we only consider the Gompertz regime, *i.e. dh*(*t*)/*dt* = *bh*(*t*) (eq3), combining with (eq1) (eq2) and differentiating on both sides, we have *d*^2^*U*/*dt*^2^ + *bdU*/*dt* = 0 (eq3) and *dU*(*t*)/*dt* = −*R h*(0), where *b* is the Gompertz ageing rate and R balance out the choice of t0 which should not alter the dynamics of (eq3). This framework could also be extended to growing populations by considering the equation *d*^2^*U*/*dt*^2^ + *b dU*/*dt* + c(*U*) = 0, where c(*U*) is zero in low levels of *U* such as in growth arrest conditions, but theoretically could be larger than zero, pushing the physiology away from senescence but toward rejuvenation.

Despite the simplicity of bacteria and its vast differences from animals, we find that concepts in evolutionary theories of ageing, such as antagonistic pleiotropy^5^, still apply. RpoS mediates trade-offs between growth and maintenance, between assimilation of nutrients and dissimilation of biomass. Sigma competition, a well-understood mechanism in the physiology of bacterial stress response, provides a molecular basis for such trade-offs often only hypothesised in metazoan organisms. Activated by nutritional deprivation and shown to decrease the ageing rate, GSR has immediate analogies in calorie-restriction-induced longevity of metazoan model organisms. Whereas in many studies on animal models, calorie-restriction-related pathways have only been shown to promote longevity, our demographic data allow us to show that GSR indeed changes the rate of ageing. It can be thought of as not only providing resistance against stress, but also insurance against prolonged growth arrests, or in other words, protection against the progression of time.

In the future, the experimental approach and statistical framework described here could combine with classical genetic methods to examine further ways hazard dynamics could be perturbed, and to reveal dynamic features of the complex network shaping cellular senescence under various environmental constraints. Future modelling efforts should clarify how adaptive processes could optimise redundancy of living systems, in trade-offs with other traits, to produce the Gompertz law.

## Acknowledgements

This study was supported by the Axa Research Fund Chair on Longevity. We thank X. Song for contributions on image analysis, D. Misevic and other member of INSERM U1001 for their advice and discussions.

## Author contributions

Conceived and designed project: Y.Y., L.X., F.T., A.B.L.; Conceived and designed experiments: Y.Y., A.B.L.; performed experiments: Y.Y. A.S. C.L.; Data analysis: Y.Y., Wrote the paper: Y.Y. with contribution from A.B.L.

## Methods

### Experimental methods

#### Microfluidic chip fabrication

Our Polydimethylsiloxane (PDMS) lab-on-a-chip system consists of two layers, containing the flow channel and the array of cell-sized chambers. The two layers are fabricated separately using soft-lithography technology, and then bonded together to produce the microfluidic device (Fig. S2)

To fabricate the negative master for the piece containing the flow channel, SU8 3050 photoresist (Microchem, MA, USA) was patterned on a silicon wafer using photolithography. SU8 3050 was spin-coated on a silicon wafer at 4000 rpm for 30 seconds, baked at 95 °C for 15 minutes, and then subjected to UV exposure (25 s, 10m W/cm2). After post-exposure baking (95 °C for 5 minutes), the master was developed using SU8 developer (Microchem, USA), rinsed with isopropanol and dried with filtered nitrogen.

The master for the layer with cell-sized chambers was fabricated using reactive ion etching (R.I.E.) technology on a silicon wafer. The mask was patterned on a silicon wafer using photoresist AZ5214 (Microchem, USA), and a 100-nm layer of nickel was sputtered on to the substrate. A lift-off procedure was applied to remove the photoresist layer yielding the metal mask for the R.I.E. process. By adjusting the R.I.E. parameters (SF6 = 4 sccm, CHF3 = 16 sccm, pressure = 10 mTorr and power = 30 W), we managed to achieve a large array of micro-pillars with high aspect ratio (diameter = 1.2 μm and height = 6 μm).

To form the device, PDMS mixtures (RTV615, Momentive Performance Materials Inc., Waterford, NY) were poured (flow channel) and spin-coated (array) onto the masters to a thickness of 5 mm (main channel) and 80 μm (array) respectively. Heat curing initially formed solid PDMS layers with patterned surfaces. After drilling inlets and outlets through the flow channel layer, and mounting the array layers onto cover glasses, the two layers were then bonded together using oxygen plasma (90s, 1000 mTorr). Finally, the assemblies were cured at 80 °C overnight to produce the integrated microfluidic chips. On the day of use, the wetted surfaces of the PDMS chip were first activated by 90s exposure to oxygen plasma (90s, 250 mTorr) immediately followed by infusion of 20% (v/v) polyethylene glycol (PEG400) solution to prevent bacterial adhesion and biofilm formation.

#### Media preparation

All equipment used for media preparation, sterilisation and infusion were made of non-leaching materials (glass, Polytetrafluoroethylene or similar perfluoropolymer material) to avoid contamination with trace level carbon sources from leachable plastic additives (see supplementary material for details). Media were filter sterilised (0.2 μm) to avoid volatile organic contamination during autoclaving, and glassware was sterilised by dry heat. Carbon-free minimum media mentioned below refer to those prepared in this fashion.

#### Strain information

All lifespan distributions described in the main text are measured for strains of the Keio *E. coli* BW25113 strain (“widltype”) single-gene knockout collection^40^. For the knockout strains, the presence and location of genomic inserts were verified by kanamycin resistance and PCR amplification. The general stress response phenotypes of Δ*rpoS* and Δ*rssB* were verified using the catalase test. In addition, in developing the microfluidic device and validation of our method, we used an MG1655 derived *E. coli* strain with a chromosomally integrated CFP under P2rrnB constitutive promoter^41^ (Fig. 2b).

#### Cell culture and loading

Single isolated colonies of the bacterial strains *E. coli* wildtype, Δ*rpoS* and Δ*rssB* were grown overnight in minimal medium (1 × M9 salts, 2 mM MgSO4 and 0.1 mM CaCl2) supplemented with 20% w/v glucose (final concentration 0.4%). The following day the overnight cultures were diluted 200-fold in 50 mL of fresh medium in 250 mL Erlenmeyer flasks and grown to early exponential phase (OD600 = 0.2). This growth phase was chosen to guarantee that the variation of birth time among cells was less than one cell cycle and thus to minimise the uncertainty of lifespan measurement. Cells were concentrated by centrifugation (4,000 rpm x 15 min, 37 °C) and washed by 3 cycles of gentle re-suspension with carbon-free minimal medium and centrifugation prior to injection into the microfluidic channels. Cells were then trapped into the dead-ended wells by centrifugation at 2,000 rpm for 15 min at 37 °C with a surface density up to 6.25 × 10^4^ cells/mm^2^. The main channels were then thoroughly washed with carbon-free minimal medium.

#### Experimental setup and microscopy

A constant flow of carbon-free M9 minimal medium at 20μl per hour was provided to the micro-channels using a high-precision syringe pump (Harvard Apparatus PHD 2000 Programmable) and Hamilton GC-grade glass/PTFE syringes (Gastight 1000 Series). PTFE tubing was used to connect the syringes to the microfluidic chip. The medium was supplemented with 1.5% (v/v) polyethylene glycol (PEG400) to prevent unspecific adherence of cells to the channels and 5 μg/mL propidium iodide, as a fluorescent indicator of cell viability, was added. Cell viability was monitored using temperature-controlled (37 °C) automatic time-lapse microscopy (Zeiss AX10, 63x oil-immersion objective, controlled with MetaMorph^®^ software). Focus was maintained by a Z-scanning maximum-contrast procedure using phase-contrast illumination. Focus was re-adjusted before each imaging cycle for every position and maintained within a Z-range of 0.2μm around the maximum-contrast Z-position. Phase-contrast and fluorescence images (P.I. signal ex. 546nm/12nm, em. 605nm/75nm) were acquired for every stage position once every hour for up to 150 hours.

#### Growth phenotypes

Growth phenotypes of aforementioned *E.coli* strains on selected carbon sources in minimal media were measured in the 96-well format using a TECAN Spark^®^ microplate reader. To induce the appropriate metabolic enzymes before growth curves can be measured, strains were first grown for 24 hours in minimal medium (M9, as in those used for microfluidic experiments) supplemented with the assayed carbon sources. The optimal densities of these cultures were determined, and diluted into fresh media identical to those used for the overnight cultures. The dilution ratios were chosen so that all experimental cell cultures have an initial optimal density of OD600 = 0.002. For each experimental well on the microplates, 50μl mineral oil was added to 100μl cell culture to prevent evaporation during the experiments. The microplates were then maintained at 37°C, and shaken constantly in double-orbital motion at 150rpm by the plate reader. Microplates with flat-bottomed wells were used to maximise agitation. OD_600_ readings were taken every 10min. The growth phenotypes used in Fig. 6 are based on minimum media culture supplemented with 60mM acetate as the carbon source.

### Statistical and computational methods

#### Image analysis

The cells in our experiments were trapped in an evenly spaced 2D-grid. The fluorescence signals of every cell throughout their whole lifespans could be extracted at fixed positions, once images in the time-lapse stack were properly registered. A simple registration procedure might misidentify one cell for another because the cells are vertically imaged and look very similar to each other if only local features are considered. To register the images based on global features, such as the presence/absence of cells at individual grid positions, we devised a two-pass, coarse-to-fine registration strategy. Cells were first identified and segmented within the images using the Point-Spread Function^42^. These segmented binary image stacks containing global information were registered in a coarse pass using least-square minimisation. The obtained 2D-translations were applied to the original images. A second fine registration pass was executed on these pre-registered original images using the Pyramid Approach^43^. After registration, the salient positions of cells were detected on the Z-projected images of the whole time-lapse stacks, and fluorescence time-series were extracted from these positions.

#### Regularised estimation of P.I. signal

To determine the true P.I. signal of each individual cell and to remove the noisy effects of focusing fluctuations, we designed and implemented a correction algorithm to the raw fluorescence intensity time-series (Fig. S3). Our three-step algorithm estimated the focusing noises of each imaging position, and deduced them from the raw time-series to arrive at the true P.I. signals of each cell. The first step of our algorithm took advantage of the fact that focusing fluctuations should change synchronously the intensities of all fluorescent objects within an imaging position, while the true P.I. signal from the cells should move independently of each other. By averaging the fold changes in intensities over all cells within a given time-lapse image stack, the focusing noise was enhanced while cell-specific signals were spread over hundreds of independent time-series. In the second step, the averaged fold changes was decomposed into focusing noises and population-wide P.I. trends. This was possible because noise from focusing should have very quick fluctuations (small autocorrelation time on the order of imaging cycles) yet no long-term trends (focusing was maintained with a narrow Z-range of 0.2μm). We applied a Total Variation Regularisation algorithm^44^ to effectively de-noise the average fold changes to produce the population-wide long term trends. In the last step, the population-wide P.I. trends were combined with the cell-specific signals to recover the true P.I. signals used to determine the times of death. See Fig. S3 for the details and the effects of the algorithm.

#### Survival analysis

Populations within the same microfluidic channels were followed at multiple imaging positions, distributed evenly throughout the flow channel. We tested the statistical consistency and homogeneity of lifespan distributions among subpopulations at different imaging positions within the same channel (see Fig. 5a and Fig. S4 for examples of such subpopulations). The P.I. time-series data and lifespan distributions from each imaging position were visualised in the style of Fig. 2d (Fig. S4a). Statistically, we tested for non-parametric differences between subpopulation lifespan distributions using a two-sample Kolmogorov-Smirnov statistic sup{|Y(t)|}, *i.e*. the supremum of empirical distribution distances lY(t)| normalised to account for censorship^45^ (Fig. S4b).

After passing the consistency test, the subpopulations from the same channel were merged to form the experimental cohorts. The lifespan data of each cohort was fitted parametrically using the family of lifespan distributions described in the main text. Maximum likelihood (ML) estimators of the model parameters and their confidence intervals were obtained using the R package ‘flexsurv’. We tested the 2-, 3-or 4-parameter hazard models mentioned in the main text, with the 4-parameter Gamma-Gompertz-Makeham (GGM) model being the most general, and chose the best among these candidate models according to the Akaike Information Criterion (AIC) at their maximum-likelihood parameters.

Goodness-of-fit of the ML models were tested using the one-sample version of sup{|Y(t)|}. The best models and their fitting residues Y(t) were plotted in supplementary material. For comparison and visual inspection, we also plotted alongside the Kaplan-Meier estimators for survivorship, Nelson-Aalen estimators for cumulative hazards and their respective 95% C.I.

#### Experimental design and experimental variation analysis

Three independent experimental cohorts were tracked on separate dates for each strain. Two statistical methodologies were used to assess the level of experimental variability, and test for significant differences in lifespan distribution and its parameters among the strains. Overall variations in lifespan distribution were analysed using hierarchical clustering. The two-sample Kolmogorov-Smirnov statistic sup{|Y(t)|}^1,45^ mentioned in the previous section was used as a distance measure in complete linkage clustering. Preferential aggregation of experimental cohorts of the same genotype within the lower branches of the clustering tree was taken as evidence that lifespan distributions of the strains were significantly different.

Because the GGM model adequately described our data, we also analysed variability of its parameters and the sources of said variability. Generalised Linear Models (GLMs) with GGM as the probability distribution were built to examine the explanatory power of categorical covariates such as experimental cohorts (*exp*) or strain genotypes (*strain*). Specifically, the null hypothesis H0 that cohorts with the same genotype have the same ageing rate *b* were tested against the alternative hypothesis H1 that experiment cohorts all have significantly different ageing rates regardless of their genotypes. Since H0 is a special case of H1, both ΔAIC and likelihood ratio test were used and H1 was rejected. See supplementary material to see more details on GLM model selection and hypothesis testing.

#### Fitness models

Fitness is defined over a time period (t_1_,t_2_) as f(s,t_1_,t_2_) = Δln(*population size*)/(t_2_-t_1_), where s denotes one of the 3 strains. We calculate fitness based on time spent in “feast” or “famine” episodes. These fitness functions were extrapolated from experimental data. For “famine” episodes, changes in logarithmic population size were simply the negatives of cumulative hazards H(t_1_)-H(t_2_), so that f_famine_(*s*,t_1_,t_2_) = [H_*s*_(t_1_) - H_*s*_(t_2_)]/(t_2_-t_1_). Extrapolations from experimental data were done using the fitted GGM models. For “feast” episodes 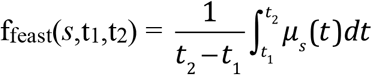. Specific growth rates *μ_s_*(*t*) and maximum growth rates 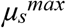 were determined from the experimental growth curves smoothed by cubic B-Splines. For “feast” episodes longer than the time at which 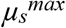 was reached, we assumed exponential growth at 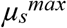.

Overall fitness f(*s*, k_1_, k_2_) of each strain at a given set of environmental parameters k_1_ and k_2_, is calculated by averaging the episodic fitness ffamine and ffeast over 5000 episodes of both “feast” τ_e,i_ and “famine” τ_m,i_, where i = 1, …, 5000. Episode lengths τ_e,i_ and τ_m,i_ are independently drawn from exponential distributions parameterised by the two ecological rates, k_1_ and k_2_, as shown in Fig. 6a. In summary, the formula for over fitness is 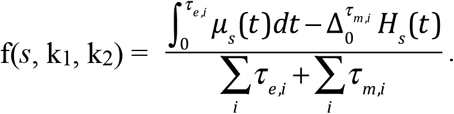.

For visual comparison in the form of the colourmap in Fig. 6b, for each set of environmental parameters (k_1_, k_2_), the fitness of the 3 strains are combined into a 3-tuple and normalised using the softmax function. If we simply denote average fitness of strain *s* as f_*s*_, the formula for the elements of 3-tuple is 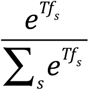, where T is the comparison timespan and is chosen to be 200 hours in Fig. 6b.

#### Programming codes and data availability

The image analysis procedure described above was implemented in Java as an ImageJ plugin. Fitness modelling, plotting and timeseries analyses including noise correction and time-of-death determination were implemented in Python. We relied on R package ‘flexsurv’ for the core algorithm of survival analysis and GLM, and Python package ‘rpy2’ was used for interoperability between the Python and R codes. All source codes and data are made available through GitHub.

## Supplemental Information

**Figure S1.**
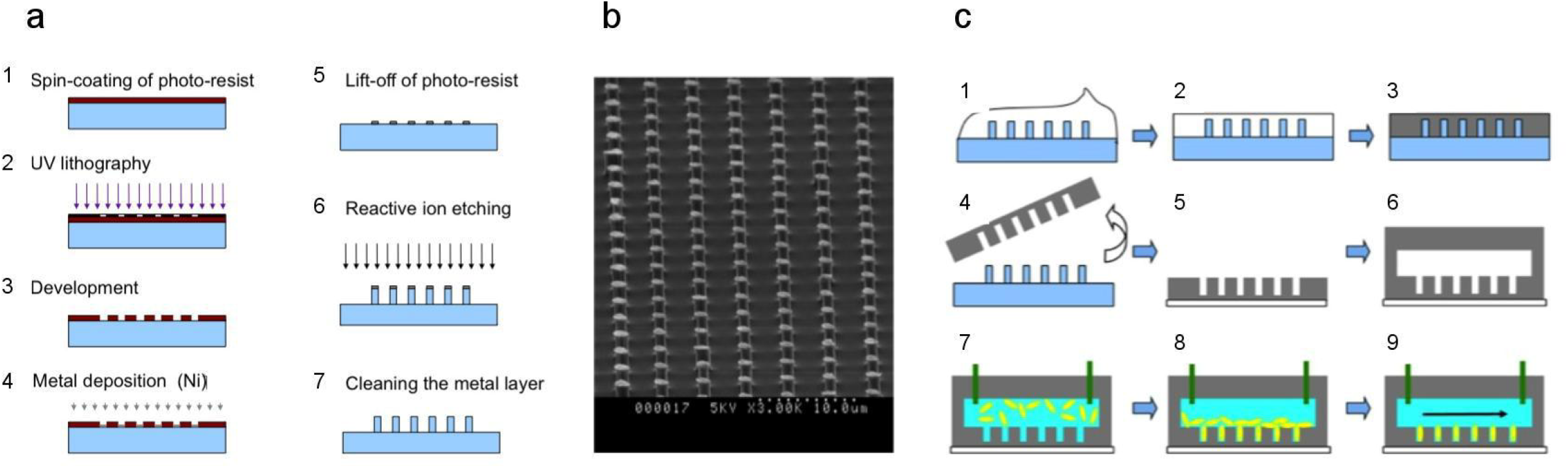
Fabrication and implementation of the 2-layer microfluidic chip. **a**, Fabrication protocol of the master for the array of cell-sized chambers. The blue shapes represent the transformations of one piece of silicon wafer. **b**, the electro-microscopy image of the silicon master. Spatial period of the array is 4 μm; the diameter and height of the pillars are 1.2 μm and height = 6 μm, respectively. **c**, The production of the 2-layer PDMS chips and the preparation and loading of the chip for lifespan tracking. White and black shapes represent uncured and cured PDMS respectively. Cyan and yellow represent cell culture media and bacterial cells.

**Figure S2.**
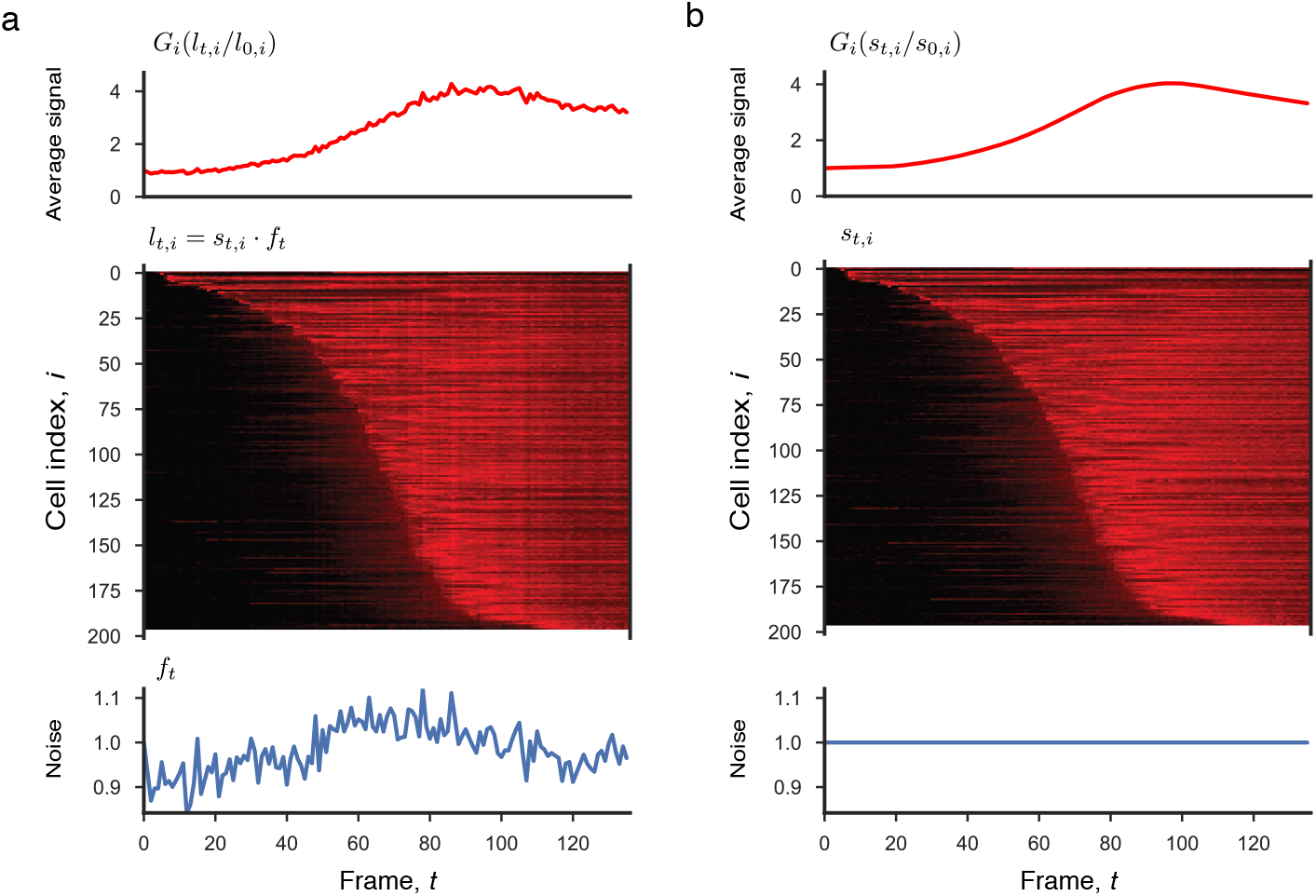
Regularised estimation of P.I. Signal to remove noise from focusing fluctuations. Average raw intensities (top), individual PI time-series from single cells ranked by estimated lifespan (middle) and estimated focusing noise (bottom) from one sample imaging position, **a**, before the algorithm **b**, after the algorithm. We assume that focusing fluctuations affect all areas within an image positions similarly, introducing a cell independent multiplicative noise *f_t_* term to the true PI signal *s_i,t_* of each cell. Thus the observed intensities extracted from the image stack are *l_i,t_* = *f_t_s_i,t_* assuming *f_t_* =1. *f_t_* can be visualised as vertical stripes in the middle panel of **a**. The first step of the algorithm takes the geometric mean of *l_i,t_*/*l_i,0_* over i, so that *G_i_*(*l_i,t_*/*l_i,0_*) = *f_t_G_i_*(*s_i,t_*/*s_i,0_*), *i.e*. the average log raw intensities consist of two terms: the focusing noise and the average PI signal fold changes. In the second step, as explained in Methods, the average PI signal *G_i_*(*s_i,t_*/*s_i,0_*) is estimated using Total Variation Regularisation^44^ of *G_i_*(*l_i,t_*/*l_i,0_*), while the remaining component is assumed to be *f_t_*, assuming that *G_i_*(*s_i,t_*/*s_i,0_*) changes gradually while *f_t_* has short temporal correlations and large temporal derivatives. *G_i_*(*s_i,t_*/*s_i,0_*) and *f_t_* of the example imaging position are visualised in the top panel of b and bottom panel of **a** respectively.

**Figure S3.**
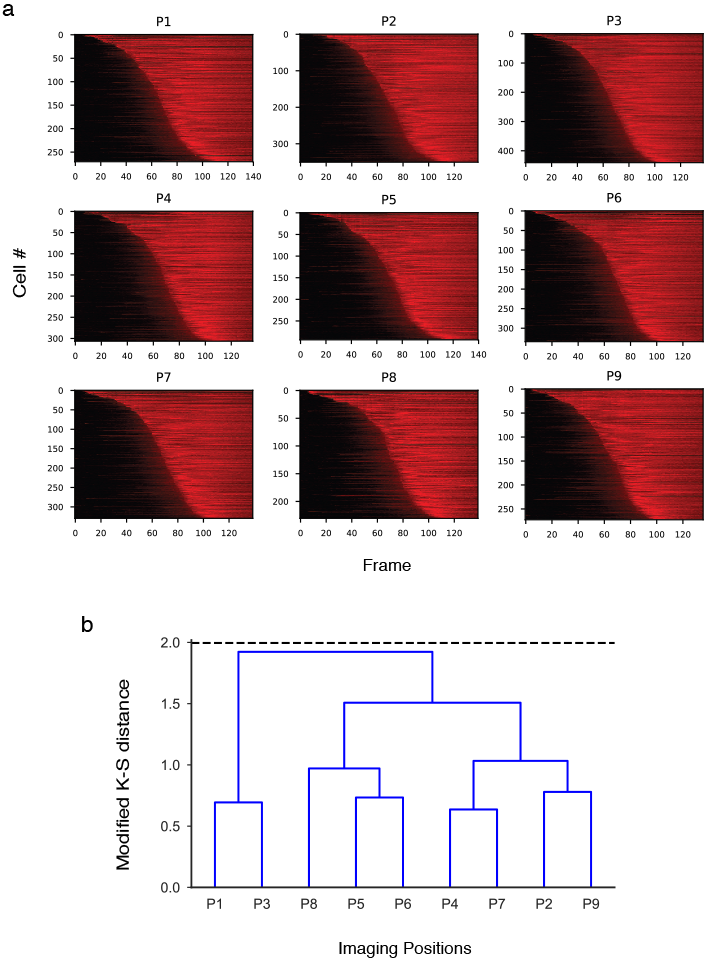
Visualisation and non-parametric statistics of variability among imaging positions within one experimental cohort. **a**, Lifespan distributions and PI time-series of sub-populations from all imaging positions within one microfluidic channels, visualised in the style of of Fig. 1d. **b**, Hierarchical clustering of these sub-populations. Standard agglomerative clustering algorithm is applied using complete linkage and the 2-sample modified Kolmogorov-Smirnov statistic sup{|Y(t)|} as a distance metric. Dashed line corresponds to 0.05 two-tailed significance level adjusted for multiple testing.

**Figure S4.**
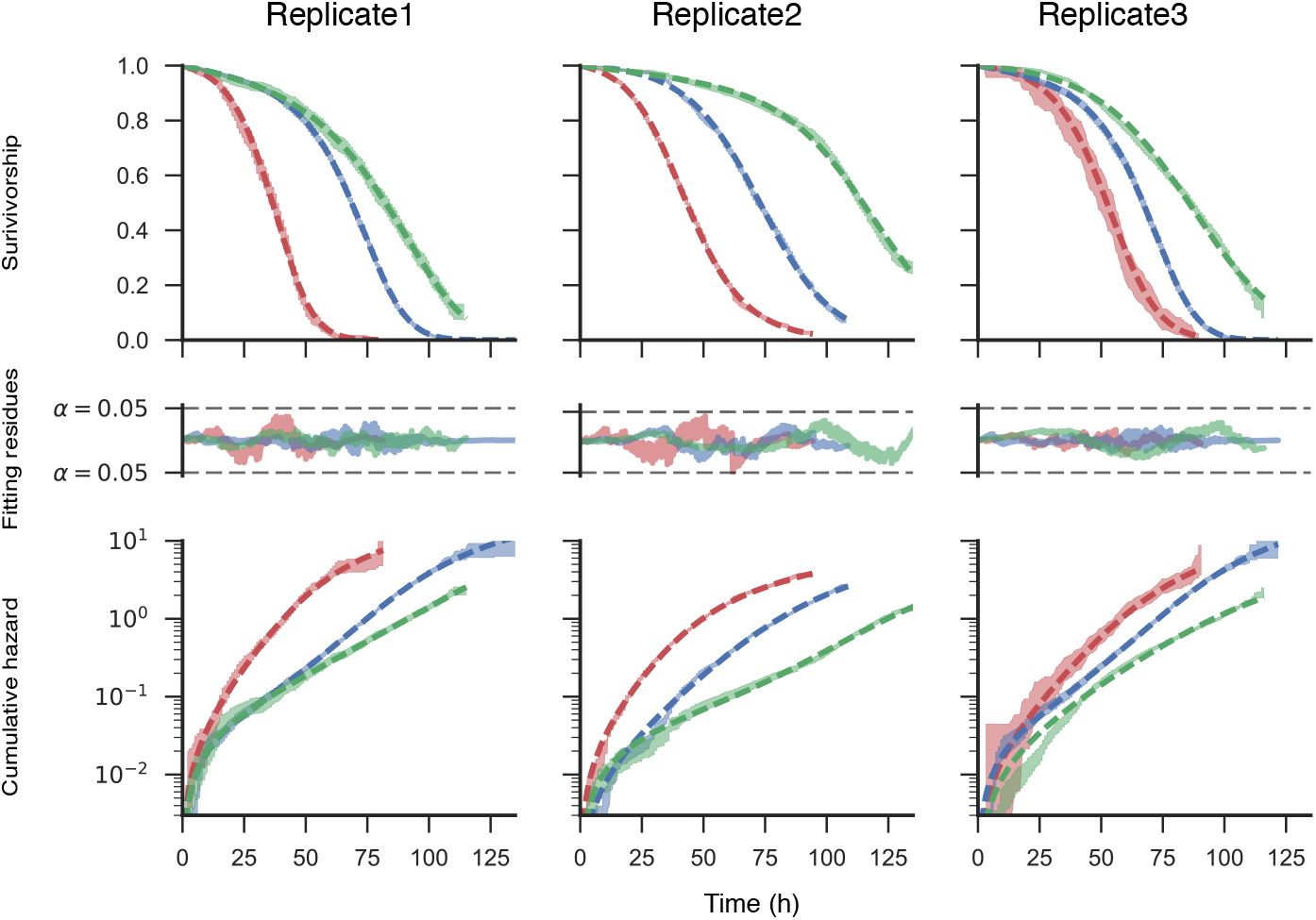
Mortality statistics and goodness-of-fit of 3 independent experimental replicates for each of the 3 strains in Fig. 2. Each column corresponds to each of the 3 experimental replicates, while each row corresponds to a different mortality statistics: top, Survivorship; middle, fitting residues; bottom, cumulative hazards. The cumulative hazards are plotted in log scale so that exponential regimes described by the Gompertz law can be visualised as straight lines. Statistics of the 3 strains are plotted using the same colour code in Fig. 2: blue, *wildtype*; green, Δ*rssB;* red, Δ*rpoS*. Coloured dashed lines in all panels are the most likely Gamma-Gompertz-Makeham model, assessed by AIC. Coloured bands in the top and bottom panels are the 95% CI of the Kaplan-Meier (survivorship) and Nelson-Aalen (cumulative hazard) estimators respectively. The fitting residues are the modified Kolmogorov-Smirnov statistic Y(t) whose extrema can be used to test the goodness-of-fit of the model in the top and bottom panels. Black dashed lines in the middle panels indicate the 0.05 significance level for sup{|Y(t)|} (H_0_: model fits the data).

